# Modeling and simulating networks of interdependent protein interactions

**DOI:** 10.1101/229435

**Authors:** Bianca K. Stöecker, Johannes Köester, Eli Zamir, Sven Rahmann

## Abstract

Protein interactions are fundamental building blocks of biochemical reaction systems underlying cellular functions. The complexity and functionality of these systems emerge not only from the protein interactions themselves but also from the dependencies between these interactions, e.g., allosteric effects, mutual exclusion or steric hindrance. Therefore, formal models for integrating and using information about such dependencies are of high interest. We present an approach for endowing protein networks with interaction dependencies using propositional logic, thereby obtaining *constrained protein interaction networks* (“constrained networks”). The construction of these networks is based on public interaction databases and known as well as text-mined interaction dependencies. We present an efficient data structure and algorithm to simulate protein complex formation in constrained networks. The efficiency of the model allows a fast simulation and enables the analysis of many proteins in large networks. Therefore, we are able to simulate perturbation effects (knockout and overexpression of single or multiple proteins, changes of protein concentrations). We illustrate how our model can be used to analyze a partially constrained human adhesome network. Comparing complex formation under known dependencies against without dependencies, we find that interaction dependencies limit the resulting complex sizes. Further we demonstrate that our model enables us to investigate how the interplay of network topology and interaction dependencies influences the propagation of perturbation effects. Our simulation software `CPINSim` (for Constrained Protein Interaction Network Simulator) is available under the MIT license at http://github.com/BiancaStoecker/cpinsimandviaBioconda (https://bioconda.github.io).

**Author summary:** Proteins are the main molecular tools of cells. They do not act individually, but rather collectively in order to peform complex cellular actions. Recent years have led to a relatively good understanding about which proteins may interact, both in general and in specific conditions, leading to the definition of *protein interaction networks.* However, the reality is more complex, and protein interactions are not independent of each other. Instead, several potential interaction partners of a specific protein may compete for the same binding domain, making all of these interactions mutually exclusive. Additionally, a binding of a protein to another one can enable or prevent their interactions with other proteins, even if those interactions are mediated by different domains. Hence, understanding how the dependencies (or constraints) of protein interactions affect the behaviour of the system is an important and timely goal, as data is now becoming available. Here we present a mathematical framework to formalize such interaction constraints and incorporate them into the simulation of protein complex formation. With our framework, we are able to better understand how perturbations of single proteins (knockout or overexpression) impact other proteins in the network.

## Introduction

A central goal in cell biology is to understand how cellular functions emerge from the collective action of interacting proteins. High-throughput protein-protein interaction detection techniques, including yeast two-hybrid and mass spectrometry [1–3], can provide static snapshots of complete interactomes, as demonstrated with several organisms [4,5]. The obtained information is typically modeled as networks, i.e. undirected graphs with nodes and edges corresponding to the proteins and their pairwise physical interactions, respectively [6–8]. However, a fundamental feature of protein networks is that the interactions between proteins are dependent on each other. A key mechanism generating interaction dependencies is allosteric regulation, in which a protein undergoes conformational change upon one interaction which affects its capability to bind other proteins [9]. Another major mechanism for interaction dependencies is mutual exclusiveness arising from steric hindrance that prevents proteins from binding simultaneously to too close or identical protein domains (Figure 1).

**Figure 1.**
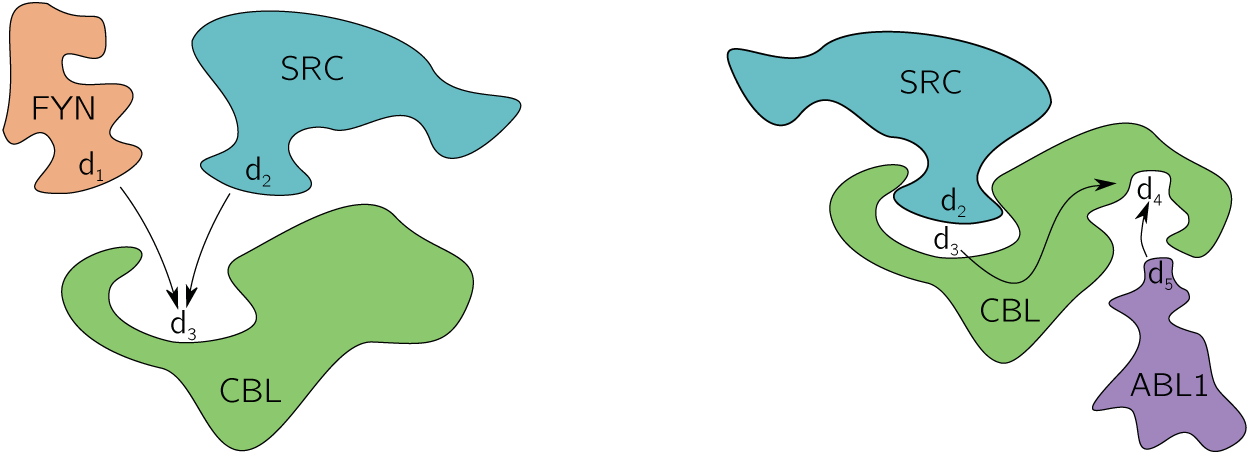
Interaction dependencies limit simultaneously possible protein interactions. Two or more proteins can compete on the same binding domain (left) leading to the constraints { (CBL, *d*_3_), (SRC, *d*_2_)} ⇒ ¬ { (CBL, *d*_3_), (FYN, *d*_1_)} and { (CBL, *d*_3_), (FYN, *d*_1_)} ⇒ ¬ { (CBL, *d*_3_), (SRC, *d*_2_)}. On the other hand, one interaction can depend on another, allosteric, interaction that induces a conformational change (right). This is represented by the constraint {(CBL, *d*_4_), (ABL1, *d*_5_)} ⇒ {(CBL, *d*_3_), (SRC, *d*_2_)}.

The dependencies between interactions have profound impact on the properties of a protein network, as they constrain the set of possible protein complexes and their assembly paths [10–12]. Moreover, interaction inter-dependencies enable perturbations of one interaction to propagate along the network and affect other interactions [11,13–15]. Therefore, considering the dependencies between protein interactions and inferring their collective impact is essential for understanding the design and function of intracellular protein networks.

An example of a bioinformatics application that benefits immediately from the incorporation of dependencies is protein complex prediction. Various approaches to infer protein complexes *in silico* at large scales exist [16]. They usually rely on detecting dense regions in the plain protein network via clustering algorithms [17–20]. Studies indicate that considering mutual exclusiveness between interactions improves the quality of such protein complex prediction [21–24]. So far, no approach appears to exist that takes arbitrary types of dependencies into account.

Ultimately, a complete quantitative biochemical description of the whole biochemical system, including the concentrations and spatial distribution of all involved proteins and the kinetic constants of their interactions is desirable [25–27]. However, despite the progress in technologies for measuring these parameters in living cells, a complete description of large intracellular biochemical systems is still beyond reach. Moreover, even given a detailed description of such a complex system, insightful simulations and modeling remain challenging with current computational technology [27,28]. Therefore this approach is fundamentally difficult even for small protein networks [29,30]. A valuable simplification of this challenge can be achieved based on the observation that mutual exclusiveness and allosteric regulations typically lead to all-or-none changes in the state of the target protein interaction, and therefore can be viewed as Boolean-logic dependencies between protein interactions. Logic-based models were previously successfully used for the analysis of signaling networks [31].

In comparison to finding interactions between proteins, identifying the dependencies between the interactions is more challenging. In order to infer mutual exclusiveness between the binding of two proteins to a third one, their minimal binding domains should be identified and found to be at least partially overlaping. However, in case of non-overlapping binding sites which are in close proximity, structural information has to be incorporated in order to determine if there is steric hindrance between the binding proteins [13,14,32,33]. Similarly, structural comparisons between proteins that are known to interact with a common protein enable to infer probabilities for mutually exclusive interactions in a protein network [34]. Advances in computational protein-protein docking enable to infer protein interactions, and hence with sufficient structural resolution it can also indicate competing interactions throughout a network [35–39].

In addition to the experimental and computational challenges to identify dependencies between protein interactions, the knowledge that accumulated about such interaction dependencies is less standartized and centralizied, in comparison to protein interactions. While partial information about interaction dependencies is available in databases, a considerable amount of experimental findings which indicate interaction dependencies are textually described in scientific publications, rather than standardized for mining. Along this line, we previously established a computational approach for high-throughput mining of protein interaction dependencies from large text corpora [11]. Finally, while all of the aforementioned methods lead gradually to accumulation of organized information about interaction dependencies in large biochemical systems, a comprehensive approach to integrate this knowledge for getting a better understanding of large biochemical systems is still required.

### Contributions

So far, no unifying model appears to exist that takes arbitrary types of protein interaction dependencies (beyond mutual exclusiveness) into account. Additionally, previous work rarely considered the concentration of proteins, although they can, in particular combined with interaction dependencies, have a significant impact on the possible complexes. Here we propose a framework and a simulation method for the evaluation of complex assembly on a large scale, for hundreds of proteins with thousands of copies. We use *propositional logic* to model interaction dependencies, and provide a flexible framework for their system-wide representation that we call a *constrained protein interaction network* (more specifically, a constrained protein domain-domain interaction network, or just *constrained network* for short). We present a computational approach to simulate constrained networks for studying steady state and response to perturbations (knockout and overexpression of single or multiple proteins, changes of protein concentrations). We show how this framework enables a fast simulation and the analysis of many proteins in large networks. We then illustrate the benefits of our model on the human adhesome network, with adjusted simulation parameters to match properties of known human protein complexes. By comparing complex formation with known dependencies against complex formation without dependencies, we show that interaction dependencies limit the resulting complex sizes and have an influence on the fraction of singleton proteins of each type. We illustrate how our model enables us to study the effects of perturbations like knockout or overexpression of proteins. Thereby, we show how the interplay of network topology and interaction dependencies guides the propagation of perturbation effects across the network.

To allow others to investigate these effects, we offer our simulation software `CPINSim` (Constrained Protein Interaction Network Simulator) under the MIT license at http://github.com/BiancaStoecker/cpinsim.

## Methods

### Constrained protein interaction networks: model

A *protein-protein interaction network* may be formalized as an undirected graph *(P, I)* with a vertex *p* ∈ *P* for each protein and an undirected edge *{p,p′}* ∈ *I* for each possible interaction. In this sense, the graph describes all potential interactions, not a concrete state of interacting proteins.

Sometimes there are several possible interactions between two proteins, which can be distinguished by different binding domains. Therefore, a more fine-grained model is helpful that considers interactions between domains of proteins.

#### Definition 1

(Domain interaction network). A *protein domain* is a pair (*p, d*)consisting of a protein name and a domain name. Two protein domains (*p*_1_, *d*_1_) and (*p*_2_, *d*_2_) belong to the same protein if *p*_1_ = *p*_2_. A *domain interaction network* is an undirected graph (*P, I)* whose nodes are protein domains (*p, d*) ∈ *P* and whose edges are domain interactions {(*p*_1_, *d*_1_), (*p*_2_, d_2_)} ∈ *I*.

A domain interaction network (*P, I*) can be projected down to a protein interaction network (*P′, I′*) by defining *P*′ := {*p* | (*p, d*) ∊ *P*{ and *I*′ := {{ *p*_1_*p*_2_} | {(*p*_1_, *d*_1_, (*p*_2_, *d*_2_)} ∈ *I*}.

We now present a method for incorporating *dependencies* between domain interactions. Our method is based on propositional logic [40].

#### Definition 2

(Propositional logic). The *propositional logic* 𝔓𝖗𝖛𝖕(*Q*) over a set *Q* (the atomic units of the logic) is the smallest set of *formulas* such that

- T (True) and ⊥ (False) are formulas.
- *q* itself is a formula for all *q* ∈ *Q*
- if *ϕ, ϕ′* are formulas, so are ¬*ϕ*, *ϕ* ∧ *ϕ′*, *ϕ* ∨ *ϕ′*, and *ϕ* ⇒ *ϕ′*. (The operators ¬, ∧, ∨, ⇒ have the usual semantics “not”, “and”, “or”, and “implies”, respectively. The implication *ϕ* ⇒ *ϕ′* is equivalent to (¬*ϕ* ∨ *ϕ′*).)

In our application, the atomic units of the logic are the interactions *I*. Thereby, the satisfiability of an interaction *i ∊ I* represents whether it is possible or not in a given state (e.g. in partially assembled complexes). We describe interaction dependencies via propositional logic formulas with a particular structure (“constraints”).

#### Definition 3

(Constraint for an interaction dependency). A *constraint* is a propositional logic formula of the form *i ⇒ ψ* with *i ∊ I* and ψ ∊ 𝔓𝖗𝖛𝖕(*I*). With `C`(*I*) ⊆ 𝔓𝖗𝖛𝖕(*I*) we denote the set of all possible constraints over *I*.

A constraint *i ⇒ ψ* restricts the satisfiability of i by the satisfiability of *ψ*. In other words: Formula *ψ*; is a necessary condition for interaction *i*.

For example, the dependency of an interaction *i* on an allosteric effect due to a scaffold interaction *j* can be formulated by the constraint *i* ⇒ *j*. Mutual exclusiveness of two interactions *i, j* ∊ *I* can be modeled by the two (equivalent) constraints *i* ⇒ ¬ *j* and *j* ⇒ ¬*i*. Figure 1 shows some examples graphically.

Using propositional logic also allows defining constraints of higher order: An interaction *i* could depend on an arbitrary scaffold interaction of a given set *j*_1_,…, *j_n_*, which is modeled by the formula *i* ⇒ (*j*_1_ ∨ …;∨ *j_n_*). For example the interaction of F-ACTIN with VCL becomes possible by either ACTN1 or TLN1 binding to VCL. This leads to the constraint

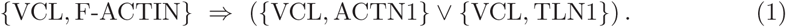

Protein domains have been omitted for readability. By combining multiple constraints, it is possible to model arbitrary combinations of allosteric effects and steric hindrance.

Now, we can define constrained protein interaction networks as a set of protein domains (nodes) connected by interactions (edges) extended by a set of constraints (dependencies between edges).

#### Definition 4

(Constrained protein domain-domain interaction network). Let (*P,I*) be a domain interaction network. Let *C* ⊆ 𝕮(*I*) be a set of constraints according to Definition 3. Then the triple (*P,I,C*) is called a *constrained protein domain-domain interaction network*, or *constrained network* for short.

### Simulation of protein complex formation

A constrained network allows us to approximate the behavior of real proteins in a cell via simulations. For a constrained network (*P, I, C*), we consider *n_p_* copies of each protein *p* to be present in the system. Together, the domains of these protein copies form a graph. Edges represent currently happening interactions between domains. In addition, we consider all domains of the same protein to be implicitly connected. Hence, the connected components of the graph represent protein complexes. Initially, each complex is a singleton protein: there are no interactions. We abstract from the spatial location of the proteins, and perform our simulation stepwise by repeatedly conducting two phases. In the association phase, each protein copy can (randomly) form new associations according to the current state, the possible interactions and the interaction constraints. In the dissociation phase, existing interactions probabilistically dissociate, potentially breaking large complexes into smaller ones. These phases are repeated until certain observable quantities reach stable levels (“convergence”, see below).

In the association phase, we iterate over all protein copies. For each copy, with a given probability *α* (association probability), a new interaction is attempted (with the complementary probability 1 − α, the protein copy will do nothing in this phase). For protein copy *p* we have Σ_*p′*_:{(_*p,d*_),_(*p′, d′*)}∊*I*_ *n_p′_* different possible interactors to choose from. To attempt a new interaction, first an interactor *p′* and then a specific domain interaction *i* = {(*p, d*), (*p′, d′*)} are randomly chosen from the possible interactions not yet established with *p*. It is then checked whether the proposed interaction i is valid, i.e., that no constraint will be violated if it is established. For this, we consider the conjunction of *i* with all present interactions *I_p_* ⊆ *I* and *I_p_* ⊆ *I* involving protein *p* or *p′* respectively and all constraints of the new interaction *i*. Consider the propositional logic formula

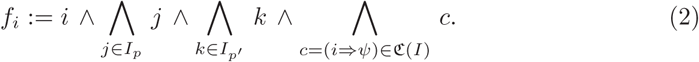

The interaction *i* can be formed if and only if *f_i_* is satisfiable. Essentially, satisfiability means that none of the constraints c contradicts the conjunction of the new and the existing interactions. Satisfiability of *f_i_* is checked (see below for an efficient algorithm), and interaction *i* is added to the simulation state in the affirmative case. If the proposed interaction is not possible in the current state, it is not added; this leads to an effective rate less than *α* for new associations.

In the dissociation phase, we iterate over each existing interaction and remove it with probability *β*. We do not check whether any constraints are violated after removal. This is motivated by the following reasoning. Consider an allosteric activation where proteins *A* and *C* can only interact if already an interaction between *A* and *B* is present. Assume a state where both interactions exist for specific copies of *A*, *B* and *C*. Now the interaction between *A* and *B* may dissociate without removing the interaction between *A* and *C*. So while that interaction is necessary for the formation of the interaction *A* and *C*, it is not necessary for maintaining it. This simplification is based on the assumption that once the allosteric activator *B* enabled the binding of protein *C* to protein *B*, the bound *C* locks the conformation of *B* in the state which is compatible for allowing this interaction. An example for such binding-mediated conformation locking is the binding of Vinculin (VCL) to Talin (TLN1), which depends on the mechanical stretching of Talin and then inhibits Talin refolding after the force is released [41].

We conduct the simulation until a steady state has been reached. Informally, this is a state where subsequent simulation steps change neither the total number of interactions (edges) in the simulation network nor the distribution of complex sizes in the network. As a proxy for the size distribution, we consider the fraction of singleton proteins (i.e., the number of non-interacting protein copies, divided by the total number of protein copies in the simulation).

As we start with no interactions, during the initial steps, the number of interactions (edges) grows until association and dissociation reach a balance where the number of interactions stabilizes. Formally, we detect this point in step s when the mean number of edges over the last ten steps (*s* – 9,…, *s*) is smaller than the previous ten-step mean (steps *s* – 10,…, *s* – 1). We then continue the simulation for another s steps to monitor the behavior and ensure that both the number of interactions and the proteins’ singleton fractions have stabilized. So the simulation runs for 2s steps when in step *s* the convergence criterion is first satisfied. Figure 2 visualizes the process.

**Figure 2.**
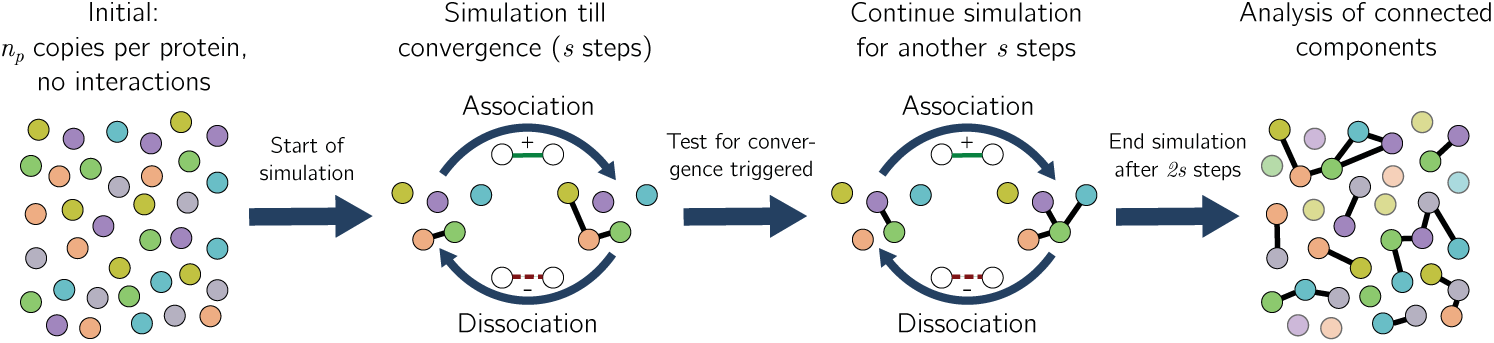
Visualisation of the simulation procedure. Starting without interactions, association and dissociation phases alternate until the convergence criterion is first satisfied after *s* steps. Then the simulation continues for another *s* steps.

### An efficient algorithm for checking constraints

We now discuss how the decision whether a proposed interaction *i* = {(*p, d*), (*p′, d′*)} does not violate any constraints can be made quickly during the simulation. This is of importance because potentially hundreds of thousands such decisions must be made during a single simulation run.

Recall that we need to evaluate whether the formula fi given by (2) is satisfiable, where *I_p_,I_p′_* in (2) are the sets of existing interactions involving the same protein copies *p, p′* as in *i*. Since *i* and all active interactions *j* and *k* have to be present, we can omit the first half of the formula and simplify the last part to ∧_*c*=(*i*⇒ψ)∊*C*_ ψ. Note that most of the ψ will consist of a single literal (e.g., a negated interaction in case of mutual exclusion). Only in the case of higher order constraints (see Eq. (1)), a disjunction remains after the simplification. These should be rare in practice. Now, we precompute the equivalent *disjunctive normal form* (DNF). A logic formula is in disjunctive normal form if and only if it is a disjunction of clauses, where each clause is a conjunction of one or more literals [40]. In other words, we transform the constraints into the form

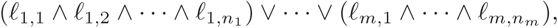

where each *ℓ.,.* is an interaction *j* or a negated interaction *¬j*. Each clause of the DNF then represents one conjunction of interactions that have to be present or absent (if occurring negated) in order for interaction i to be possible. If a clause evaluates to true when setting the already present interactions *I_p_* and *I_p′_* to true and all other interactions to false, we know that the formula *f_i_* is satisfiable. In theory, the conversion to DNF could lead to an exponential growth of the number of clauses, but as shown above, we expect most constraints to be simple, consisting of a single literal. Hence, the DNF can be calculated as follows. First, all single literals are combined into a conjunction ϕ. Second, for the first disjunction *l*_1_ ∨ *l*_2_ ∨ …, we spawn a conjunction *ϕ* ∧ *l_i_* for each literal *l_i_* and go on recursively with the next disjunction. Once the recursion is completed, we have Π_(*t*⇒ψ)∊*C*_ |*ψ*| clauses where |*ψ*| is the number of literals in the disjunction *ψ*.

Consider the subnetwork in Figure 3 with a current simulation state where one copy of protein *A* is already interacting with a copy of protein *D* and the questioned interaction is *i* = {(*A, d*_1_), (*B, d*_4_)}. The full formula from above is

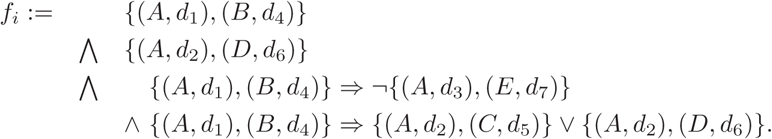

The two bottom rows represent the constraints and can be transformed into the equivalent DNF

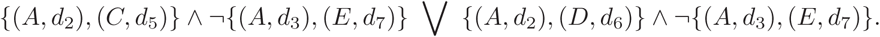

If one of the clauses is satisfied given the proposed and the existing interactions (first two rows in the formula above), then the proposed interaction is possible and does not violate any constraints.

**Figure 3.**
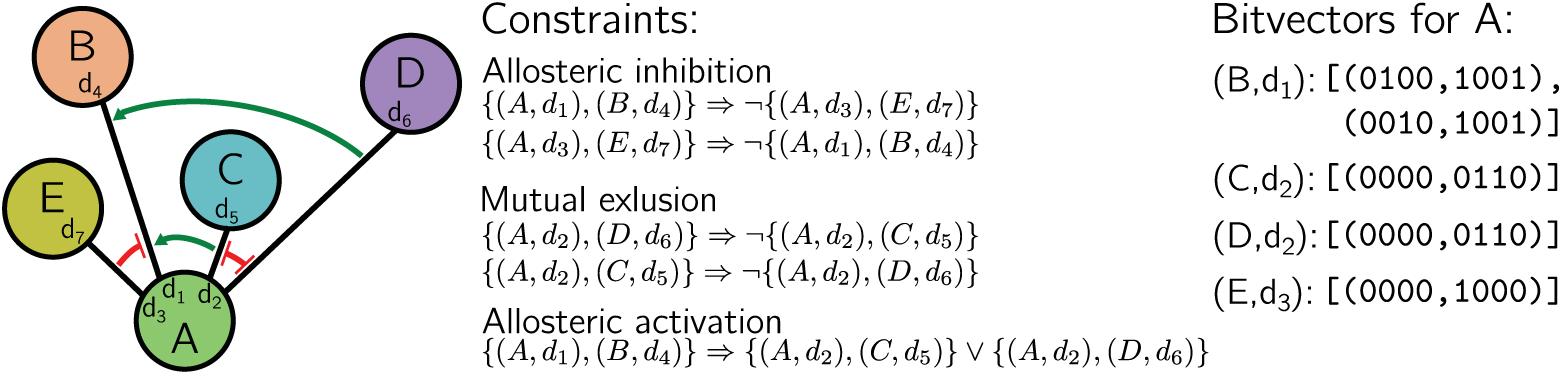
Example for the relationship between network, constraints and bit vector representation. Left: Subnetwork for Host-protein *A* with its interaction dependencies. The interaction with *B* has two independent allosteric activators *C* and *D*, that are competing for the same domain at *A*. Further, *E* is an allosteric inhibitor for the interaction between *A* and *B*. Middle: Constraints resulting from the interaction dependencies. Right: Bit vector representations of the constraints for protein *A*. The indices are assigned in lexicographical order. Since the interaction with *B* has two possible interactors, there are two clauses in the DNF and thus two pairs of bit vectors.

For each possible interaction in the system, there is one such DNF which has to be evaluated fast. For this, we propose the following approach. We first observe that each protein has a limited number of possible binding partners (Figure 6). This limits the size of the DNF clauses. For each protein, we encode the DNFs of the possible interactions using bit vectors. This representation does not change during the simulation and is shared by all copies of a protein. In addition, for each *copy p* of a protein, we represent the state of currently active interactions *I_p_* ⊆ *I* in another bit vector. This bit vector is updated whenever p enters or leaves an interaction. For a potential interaction *i* = {(*p, d*), (*p′, d′*)}, the satisfiability of f_i_ can then be efficiently checked by evaluating the bit vector representations of the corresponding DNFs for both *p* and *p′*.

**Figure 6.**
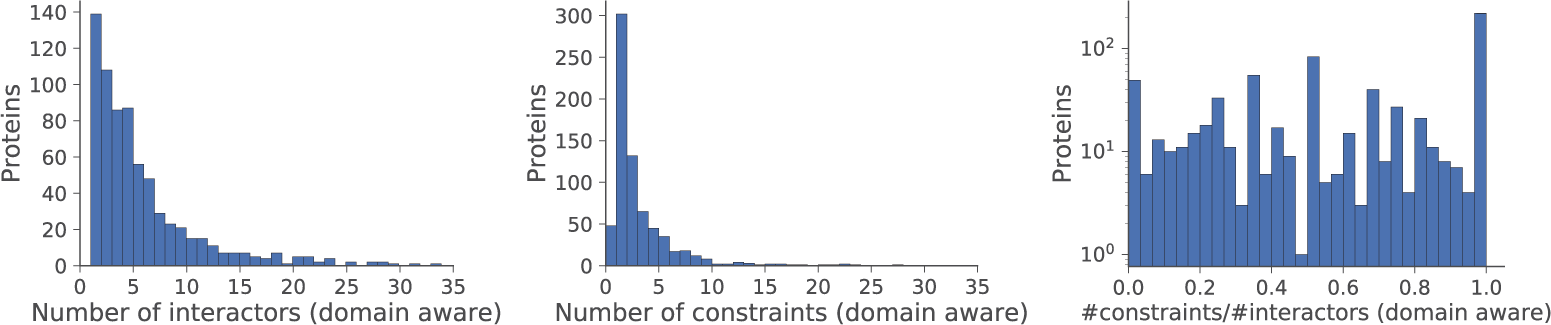
Statistics of the proteins in the constructed constrained network. Domain aware means that interactors and constraints are counted per domain. Left: Frequencies of the number of interactors; there are 17 outliers with more than 35 interactors: 38, 44, 47, 47, 47, 55, 87, 103, 118, 119, 123, 127, 135, 166, 178, 182, 242. Middle: Frequencies of the number of constraints; there are 12 outliers with more than 35 constraints: 42, 72, 73, 95, 102, 103, 106, 114, 122, 153, 166, 168. Right: Histogram of ratios of the number of constraints over the number of interactors. There are 219 proteins with a ratio of 1.

In the following we present the details of the representation. We enumerate the interactions of a protein in a convenient order and assume that the index *k_j_* of an interaction *j* can be obtained in constant time. Then, for each potential interaction of a protein, we represent each clause of the corresponding DNF by two bit vectors *b*^+^ and *b*^−^. In bit vector *b*^+^, we store the positive literals: we set the *k_j−_*th bit to one if interaction *j* occurs in a positive literal. The bit vector *b^−^* stores the negative literals by setting the *k_j−_*th bit to one if interaction *j* occurs in a negative literal. The state of currently present interactions of protein copy p is represented by a bit vector *b*^*^ with the *k_j−_*th bit set to one for each present interaction *j* ∊ *I_p_*. (The same is additionally done with another bit vector for *Ip*′.) Then, the satisfiability of the DNF can be calculated by iterating over the clauses and checking each clause against the status vector. If *b^+^* & *b*^*^ = *b*^+^ and *b*^−^ & ¬*b*^*^ = *b*^−^ (with *¬b*\* being the bitwise negation of *b^*^* and & the bitwise conjunction), we know that one clause evaluates to true. Once the iteration reaches the first satisfiable clause, we can stop, knowing that the DNF is satisfiable.

In our example we assign the indices lexicographically and get (0010,1001) and (0100,1001) as bit vectors (*b*^+^, *b*^−^) for the two clauses in the DNF. In both vector pairs the least significant (rightmost) bit reprensenting the interaction with *B* is set in the vector b^−^. This is to ensure that *A* and *B* are not already interacting with each other. The other set bit in *b*^−^ is for the allosteric inhibition of *E*. The two different set bits in *b*^+^ represent the independent allosteric activators *C* and *D*. The bit vector for the current state of active interactions is *b*^*^; = (0100). In the example, we have

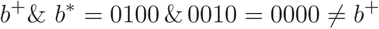

for the first clause. In contrast, the second clause is satisfiable with

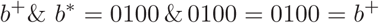

and

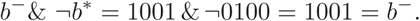

Hence, the constraints are not violated and the example interaction is possible.

### Simulation of perturbations

With a constrained network (*P, I, C*), we may simulate not only the given network, but also perturbations of it. Typical perturbations are protein *knockout* and *overexpression*. Recall that our model considers a copy number (or expression) *n_p_* for each protein *p*. Assuming that these *n_p_* copies represent a typical state of the constrained network, we can simulate a perturbation by modifying the expression of a particular protein p with a factor, i.e. 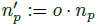. An overexpression is equivalent to a factor greater than 1, whereas a knockout corresponds to a factor less than 1. A factor o = 0 describes a perfect knockout, where no copies of the protein are left. It is of course possible to combine overexpression or knockout of different proteins in the same simulation run.

### Construction of constrained protein interaction networks

Often, it is of interest to study a certain subnetwork, that is characterized by a set of proteins *P_0_*. At the boundaries of such a subnetwork, there will be interactions that are constrained by proteins that are not part of *P_0_*. Not considering such constraints would lead to biased results. In the following, we provide a solution that includes outside proteins such that these constraints are considered as well. Given an initial set *P_0_* of proteins, protein domain interactions *I_0_* (that may involve additional proteins not contained in *P_0_*) and a set *C*_0_ of constraints over *I_0_*, we construct a constrained network (*P, I, C*) as follows.

First, note that a non-trivial constraint *c = (i ⇒ ψ*) involves at least three proteins, two in interaction i and at least an additional one in ψ. Let *P(c*) be the (multi)set of proteins mentioned in constraint *c*.

1. Select all proteins from *P_0_*.
2. Select the subset *C*_1_ ⊂ *C*_0_ of constraints that mentions at least two proteins from the initial protein set *P*_0_, i.e. *C*_1_ := {*c*: |*P*(*c*) ∩ *P*_0_| ≥ 2}.
3. Select all proteins mentioned in *C*_1_; i.e., define *P*_1_ := ∪_*c*∊*C*_1__ *P*(*c*).
4. To extend the currently selected protein set, consider constraints *C*_2_ ⊂ *C*_0_ that have an influence on the previous selected proteins; i.e.
*C*_2_ := {*c*: |*P*(*c*) ∩ (*P*_0_ ∪ *P*_1_)| ≥ 2} and select proteins *P*_2_ := ∪_*c*∊*C*_1__ *P*(*c*).
5. Define proteins *P* := *P*_0_ ∪ *P*_1_ ∪ *P*_2_, interactions *I* := {{(*p*_1_, *d*_1_), (*p*_2_, *d*_2_)} | (*p_i_, d_i_*) ∊ *P*, {(*p*_1_,*d*_1_), (*p*_2_, *d*_2_)} ∊ *I*_0_}, and constraints *C* := *C*_1_ ∪ *C*_2_.

An example for this procedure, based on an initial small subset of the human adhesome network, is shown in Figure 4.

**Figure 4.**
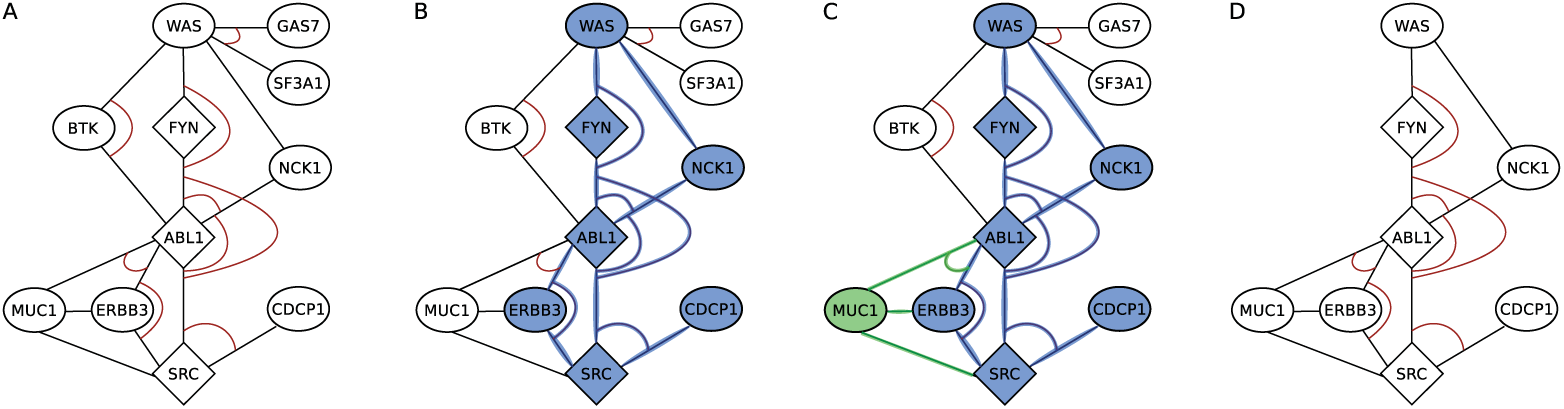
Example of the construction process of a constrained protein interaction network, starting with an initial protein set *P_0_*. We seek to find a minimal network encapsulating *P_0_* while not discarding interesting constraints. **A:** A section of the complete constrained network. The diamond-shaped nodes (◊) are proteins from the set *P_0_* (human adhesome network), the others (◯) are proteins not from *P_0_*. Black lines are interactions, red arcs indicate constraints between the interactions. **B:** Selection (blue) of all constraints and corresponding proteins and interactions, where at least two proteins are from P_0_. C: Selection (green) of proteins whose constraints have an influence on the previously selected proteins. **D**: Final set of proteins, interactions and constraints for simulation.

It remains to discuss how and where to obtain information on interactions and interaction dependencies, *I_0_* and *C_0_* in the terminology above. While protein interactions are available from various databases [42–44], information about interaction dependencies is not yet collected systematically for various reasons discussed in the Introduction. In the past, we have had success with a semi-automated text mining approach on the human adhesome network [11]. Further, competitions on binding domains can be inferred from domain interaction databases, such as DOMINO [43]. In those databases each protein interaction is annotated with a binding domain on each protein, i.e., an interval of positions in the amino acid sequence, such as the interval [540, 906]. We assume that two proteins compete for the same domain if the domains of the interactions are overlapping each other. If we have a competition between proteins without domain annotations (e.g., obtained by text mining), but each involved protein has a unique domain involved in interactions in the dataset, we assume that the constraint involves the same domains. If we cannot infer the domain in this way, we create artificial unique domains. For allosteric effects we assume that interactor and activator/inhibitor each bind to a different domain of a host protein, while competitors are all assigned to the same domain of the host.

## Results

We first describe the used adhesome interaction network and some of its statistics, as well as the chosen simulation parameters. Then we present our results regarding the running time and convergence of the simulation. Finally, we discuss the effect of constraints on complex formation and demonstrate the propagation of perturbations in the constrained network in contrast to the unconstrained network.

### Construction of the constrained extended integrin adhesome network

Since not many interaction dependencies are known, we selected a network with a high density of known constraints. In previous work, we discovered 71 interaction dependencies for the human adhesome network by systematically mining a collection of over 50000 full-text articles [11], where we searched for dependencies with at least one involved protein from the adhesome. Further we inferred competitions on binding domains from the domain interaction database DOMINO [43].

We started with the human adhesome proteins as initial set *P_0_* in the construction described above. In this initial network there are 121 proteins and 392 interactions (between only these proteins) as well as 139 competitions and 2 allosteric effects (resulting from text mining and DOMINO). The interactions between all selected proteins were taken from the HINT database (only binary interactions, [44]). Applying the described network construction leads to a network with 718 proteins (table 1),Figure 5).

**Table 1.** Characteristics of the constrained protein interaction network used for simulation (human adhesome network): Row “initial” refers to the network consisting of only nodes from the adhesome (*P_0_*). Row “extended” describes the complete constrained network constructed incrementally from *P_0_*, as described in Methods.

| network | proteins | interactions | competitions | allosteric effects |
| --- | --- | --- | --- | --- |
| initial | 121 | 392 | 139 | 2 |
| extended | 718 | 2933 | 2753 | 21 |

**Figure 5.**
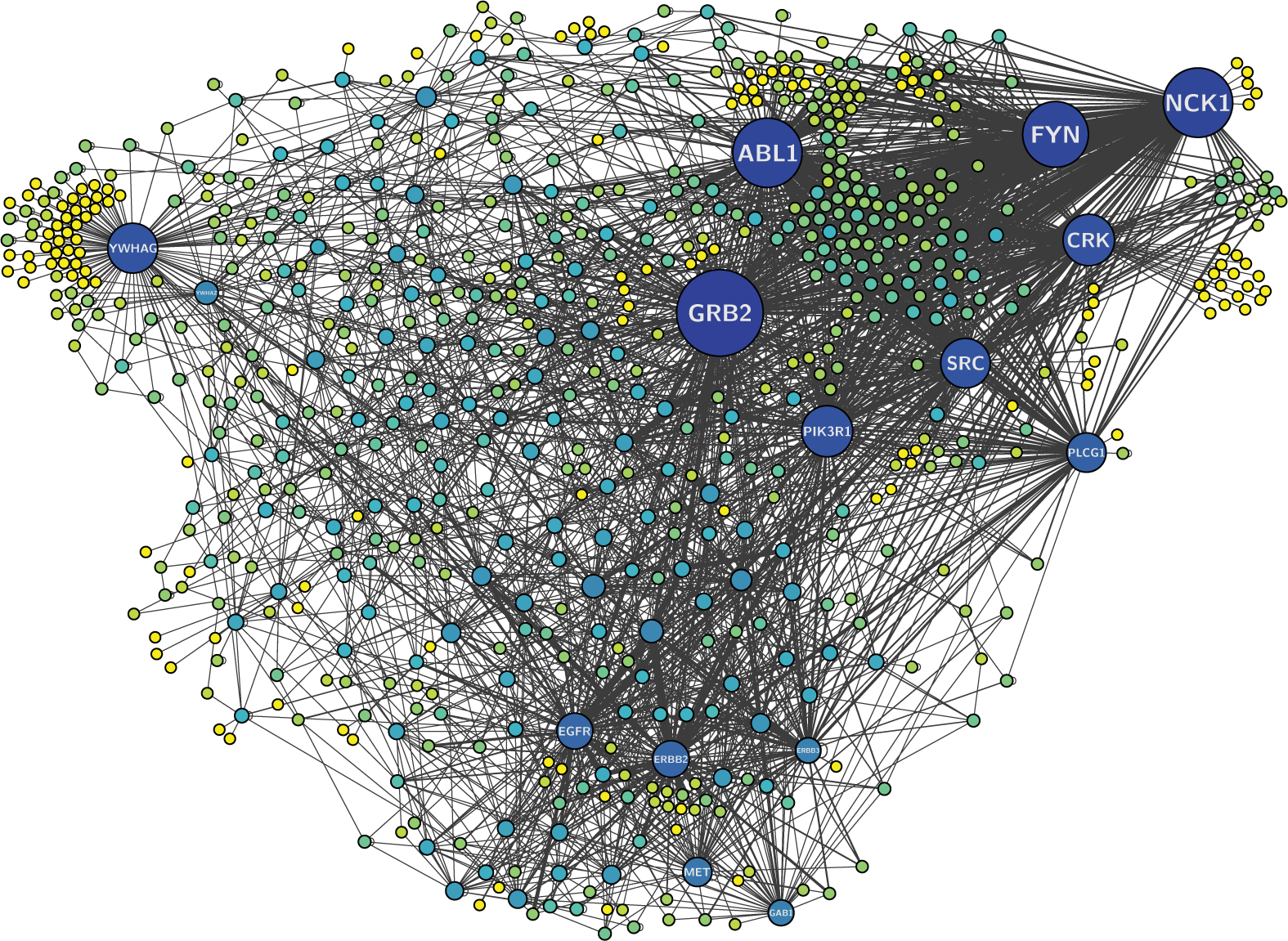
Constructed protein network (constraints not shown). Node color and size represents the number of interactors. Yellow nodes have a single interactor; the number of interactors increases with the amount of blue. Key proteins with many interactors are shown by name.

In the resulting network, 139 proteins have only one domain and one interactor, while most of the proteins have between two and six interactors. Interactors are counted with regard to the domains, meaning that the same protein is counted as two interactors if the interactions are at different domains. The distribution of the number of interactors is shown in Figure 6 (left).

There are 50 proteins that are not part of a constraint. Most proteins are part of one constraint, while some proteins are part of over one hundred constraints. The distribution of the number of constraints is shown in Figure 6 (middle).

### Choice of simulation parameters for the extended human adhesome network

Given the constrained network, the main parameters to adjust are the association probability *α* and the dissociation probability *β* (see Methods, Simulation of protein complex formation). The choice is guided by two criteria. First, the resulting complex size distribution should approximately reproduce known complex size distributions. Especially, we want to avoid the formation of overly large unrealistic complexes (several hundreds to thousands of proteins). Second, as the simulation consists of two discrete steps (association and dissociation), it has to be sufficiently fine-grained that the complex size distributions after association and dissociation phase do not differ significantly in the steady state. This excludes large probabilities.

We systematically evaluated different combinations of *α* and *β* and found that the reasonable parameter space is restricted to *α* ≤ 0.1 and *β* = ƒ. *α* with a factor f in the interval [2.0,10.0]. In this parameter range, we compared the simulated complex size distribution at steady state with the complex size distribution of known complexes. The known complexes were taken from the CORUM database provided by the Munich Information center for Protein Sequences (MIPS) [45]. The database contains manually annotated protein complexes from mammalian organisms without regard to the connections of proteins in the complexes or the multiplicity of proteins.

Figure 7 shows a histogram of complex sizes for human complexes in the CORUM database (left) and the complementary cumulative distribution functions (ccdf) of CORUM complexes and simulated complexes for a particular parameter set (*α* = 0.005, *β* = 0.0125; right). For complexes with more than 20 different proteins, information is sparse, and there are only one or two known complexes of each of those sizes. This can be explained by the difficulty of experimentally finding big complexes and represents a bias in the distribution. It can be assumed that the distribution is more accurate for the smaller complexes and thus the consistency between the known and simulated complexes for complexes with size below 20 is an indicator for a good parameter combination. The shown distribution (*α* = 0.005, *β* = 0.0125 = 2.5α) was the best fitting parameter set. Therefore those parameters were used for all following evaluations. If not stated otherwise, we consider *n_p_* = 1000 copies for each protein.

**Figure 7.**
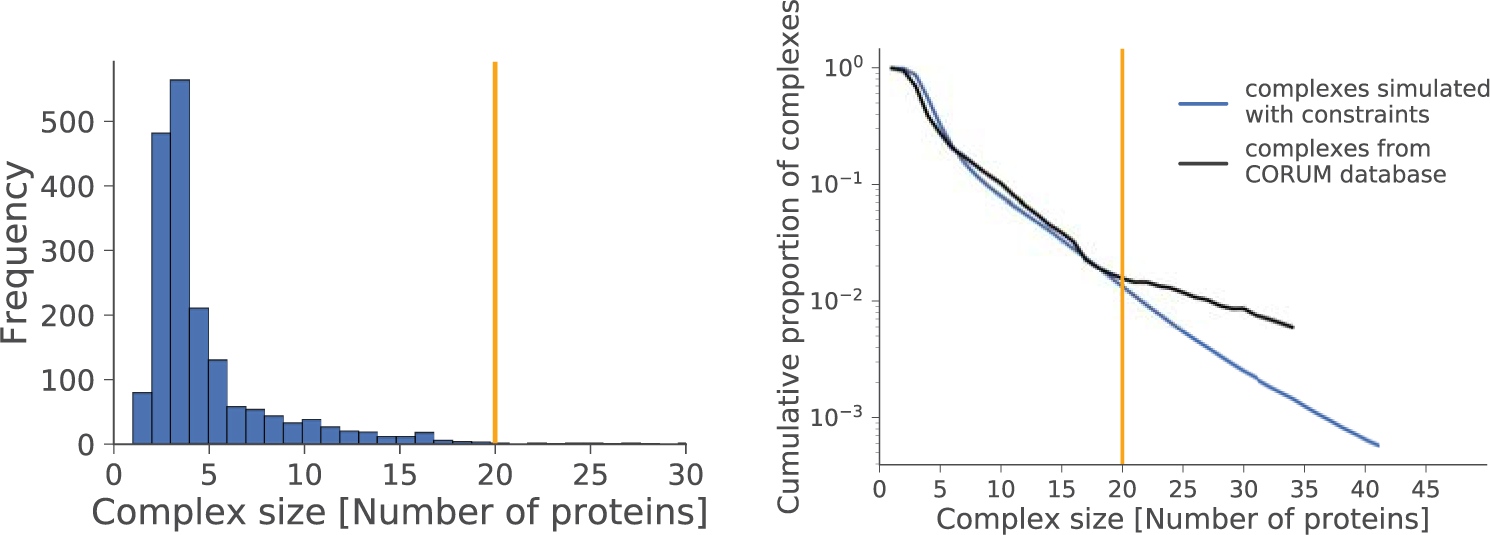
Left: Histogram of complex sizes for human complexes in CORUM database. There are 14 outliers beyond 30: 31, 32, 33, 34, 36, 37, 44, 47, 48, 78, 80, 81, 104, 143. Complex sizes larger than 20 (yellow line) occured only once or twice. Right: Comparison of the complementary cumulative distribution function (ccdf) of simulated complex sizes with constraints for *α* = 0.005, *β* = 0.0125 (averaged over 50 runs) against the distribution from the CORUM database. Cumulative distributions are capped at complex sizes where the absolute complex frequency drops below 10. Complex sizes above 20 (yellow line) occur only once or twice.

Next, we examined how many simulation steps were needed till convergence (steady state). For the chosen parameter set, between 250 and 350 steps were required (Figure 8). The convergence criterion is based on the edge density (number of interactions) in simulated complexes, which remains stable after meeting the criterion, together with the number of unbound proteins (singletons).

**Figure 8.**
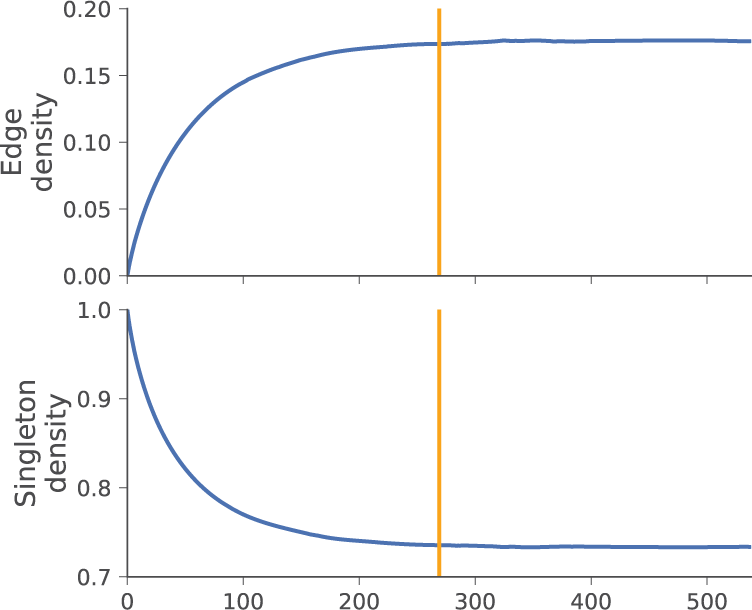
Steady-state statistics of one simulation run (with constraints); the convergence criterion is satisfied after 269 steps (yellow line), and the simulation continues for the same number of steps. Other simulations ran for comparable numbers of steps. Top: edge density (number of interactions divided by total number of protein copies in the simulation). Bottom: singleton fraction (number of singleton proteins divided by total number of protein copies).

### Running times and reproducibility of simulations

In principle, the number of proteins, interactions and constraints in a simulation is not limited, except by the available memory and, to a lesser degree, computation time. We simulated the described network with 718 protein types and *n_p_* copies of each protein for different values of *n_p_* on a single thread of an Intel Core i7-4790K processor at 4.00GHz with the time and memory requirements shown in table 2. We see that even large copy numbers can be handled in a resaonable amount of time and with an amount of RAM that is typically available on today’s common desktop PCs.

**Table 2.** Time and memory requirements for the simulation of complex formation in the extended adhesome network with 718 protein types and different copy numbers of each protein. Numbers are given for a single run, averaged over 10 runs, on a single thread of an Intel Core i7-4790K processor at 4.00GHz.

| **copies** | **time [min:s]** | **memory [GB]** |
| --- | --- | --- |
| 1000 | 04:00 | 1.23 |
| 2000 | 08:30 | 2.33 |
| 3000 | 15:03 | 2.64 |
| 4000 | 21:02 | 3.80 |
| 5000 | 26:44 | 4.49 |
| 6000 | 32:05 | 5.16 |
| 7000 | 36:58 | 6.31 |
| 8000 | 44:27 | 7.49 |
| 9000 | 48:20 | 8.17 |

We assessed whether the simulations generated reproducible results, both with and without constraints. For this, we compared the complex abundances for different runs against each other. We abstracted from network topology and multiplicity of proteins within complexes, and only considered the sets of contained proteins. Figure 9 shows the abundances of different complexes in different simulation runs, grouped by complex size. Apart from singletons, most complexes did not occur often. Yet, complexes that occurred in multiple runs tended to occur with similar frequency. Both with and without constraints, our simulation produced reproducible complexes.

**Figure 9.**
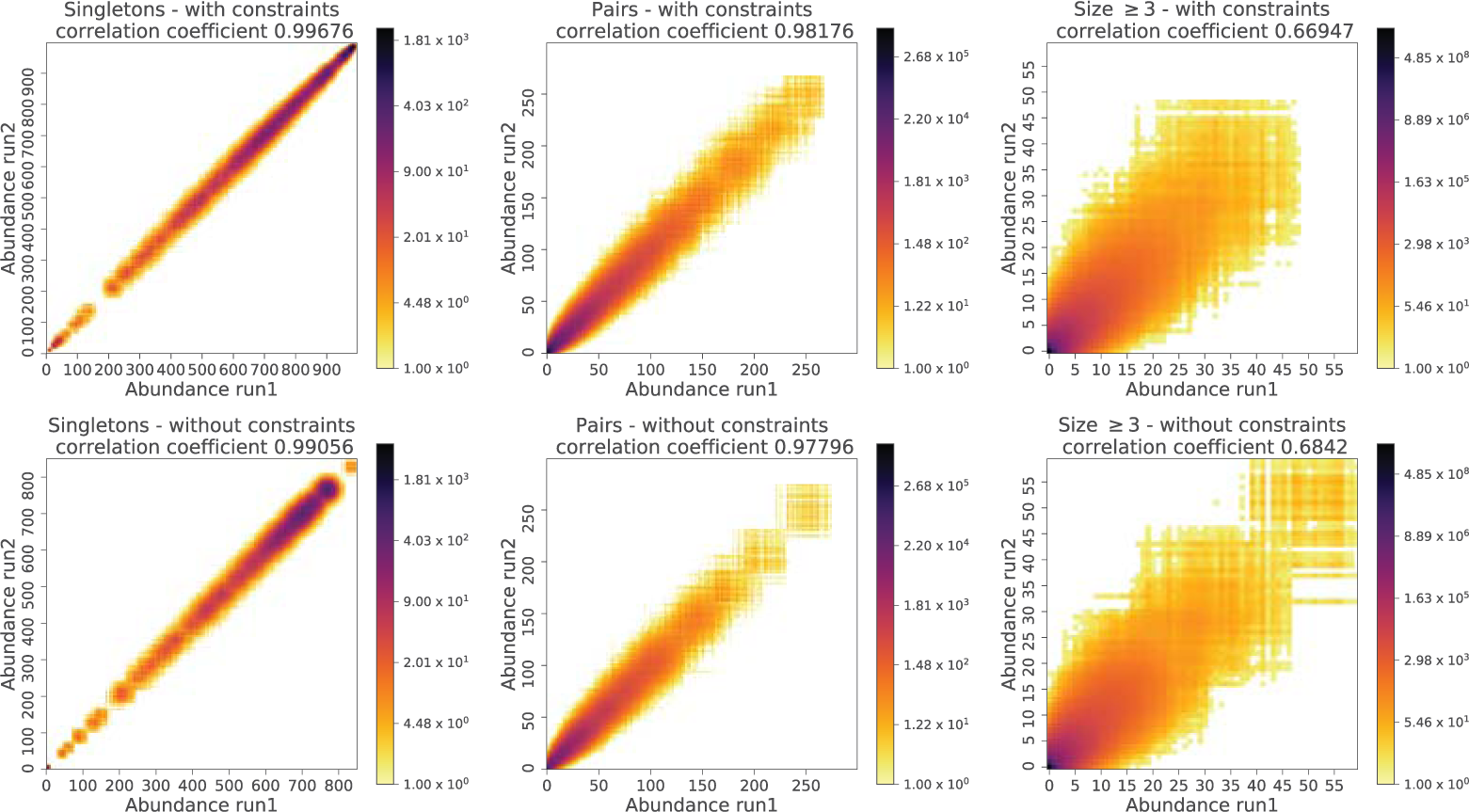
Density plot of pairwise abundances of complexes over two runs. Abundances are accumulated over the 4950 = (100.99)/2 unordered pairs of runs for 100 simulation runs, both with (top row) and without constraints (bottom tow). For this comparison, complexes are considered equal if their protein sets are equal (disregarding protein multiplicities and interactions). Note the different scales; large complexes occur less frequently than small complexes or singletons.

### Complex sizes with and without constraints

To evaluate which effect the interaction dependencies have on the simulation, we compared the simulation results with and without constraints.

For each set of constraints we did 100 simulation runs with the chosen parameters. The complex size distributions at steady state are compared in Figure 10. As could be expected, simulations without constraints lead to significantly larger complexes than simulations with them.

**Figure 10.**
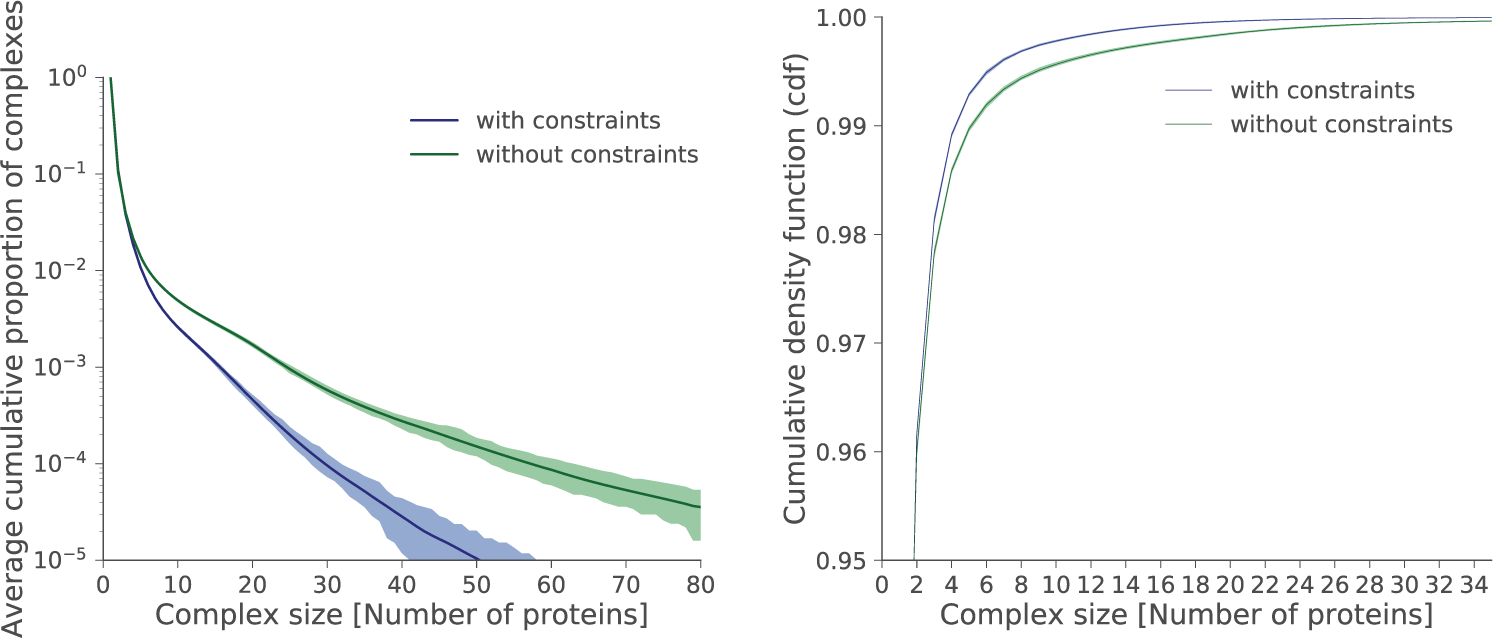
Left: Complementary cumulative distribution functions (ccdf, log scale) of protein complex sizes at steady state for 100 simulation runs with constraints (blue) and without constraints (green). The bold line depicts the mean; the shaded area depicts minimum and maximum. Right: Cumulative distribution function (cdf, linear scale) of the same runs for complex sizes ≤ 35.

The maximum complex size is 75 (averaged over runs) with constraints. All simulations without constraints develop one large complex accumulating nearly all different protein types in the network. This complex has an average size of 84 182 and no biological relevance. Further, the simulations without constraints have fewer singleton proteins at steady state than the simulations with constraints.

### Characterization of perturbation effects

In order to illustrate the capability of our framework to estimate effects of perturbations, we chose three proteins with different roles in the network, i.e. different numbers of interactors and constraints (table 3): CRK, YWHAG and ABAT. Both CRK and YWHAG have a large number of interactors and interactions, but they differ in the number of domains and additionally in their role in the network (Figure 5): CRK’s interactors include several proteins which have themselves many interactors, while YWHAG’s interactors frequently have no other interactors than YWHAG itself. In contrast, ABAT is at the periphery of the network with a single interactor.

**Table 2.**
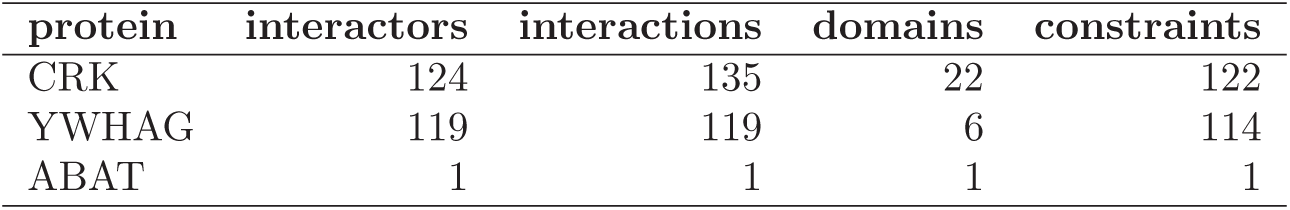
Proteins chosen for perturbation simulations (5-fold overexpression and complete knockout). For each protein, we list its number of interactors (proteins with at least one interaction with the given protein), number of model domains (including artificial unique domains, see “Construction of constrained protein interaction networks” above), number of interactions (at least as high as the number of interactors), and the number of constraints in which the protein participates.

We simulated 50 runs for each of the selected proteins with five-fold overexpression and with complete knockout of the protein, both with and without constraints.

#### Changes in complex sizes

We investigated how perturbations change the complex size distributions (Figure 11). As may be expected, perturbations of ABAT did not detectably influence the complex size distribution, neither with nor without constraints. A plausible explanation is that ABAT only has a single interactor, so its influence on the network is limited.

**Figure 11.**
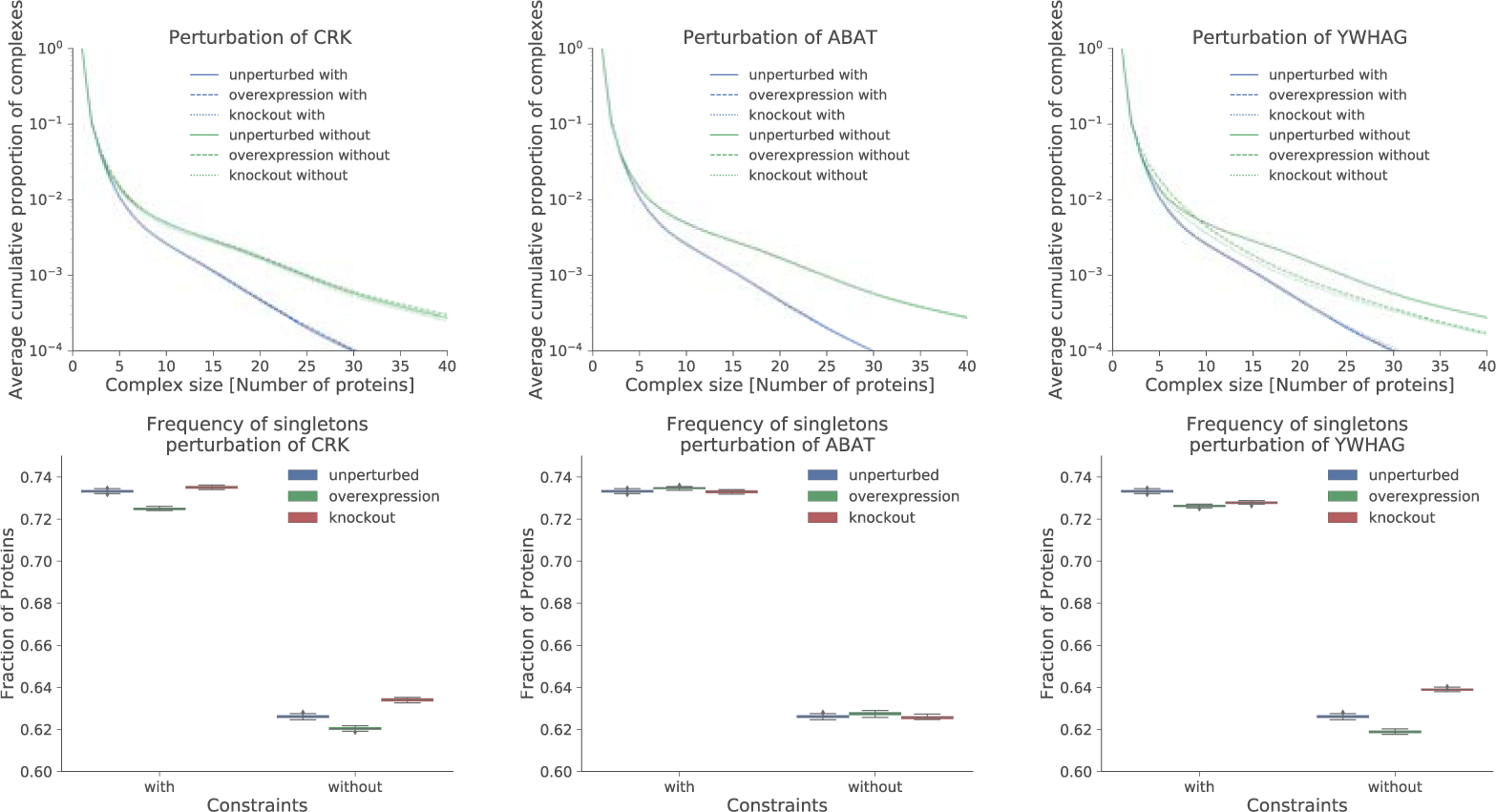
Impact of perturbations on the complex size distribution for CRK (left), ABAT (middle), YWHAG (right) under constraints (blue) and without constraints (green). Top row shows the mean complementary cumulative distribution functions (ccdf, 100 runs each with and without constraints). Solid lines depict unperturbed state, dashed lines overexpression and dotted lines knockout. The bottom row shows the numbers for singletons (unbound proteins).

Similar to ABAT perturbation of CRK had no visible global effect on larger complexes. However, singleton fractions differed with the type of perturbation (overexpression vs. knockout) and whether constraints were included in the model or not. In addition to being a central hub in the network, CRK is also a limiting factor with a small singleton fraction (i.e., most copies were bound) in the unperturbed simulations (with constraints: 0.011, without: 0.0), so a noticeable effect was to be expected. If CRK is knocked out, it can no longer act as a hub, and we observe more singleton complexes; this is true both with and without constraints. After overexpression of CRK, we observe fewer singleton complexes.

Although YWHAG and CRK are comparable concerning their number of interactors and constraints, perturbing YWHAG has different effects than perturbing CRK. With constraints, overexpression of YWHAG has little effect on larger complexes. The reason is that YWHAG does not occur in a dense region of the network, and most interaction partners can only interact with YWHAG itself while inhibiting each other, such that complex size is limited (Figure 5). On the other hand, a knockout leads to a slight increase in complex sizes. An explanation is that the few interaction partners that connected YWHAG to the rest of the network are now free to enlarge other complexes. Importantly, these effects are only visible when considering interaction dependencies. Without them, knockout leads to a major drop in complex sizes. The reason is that the family of larger complexes around YWHAG that are only possible when ignoring interaction dependencies disappears. With overexpression, there are more smaller complexes and fewer complexes of size ≥ 10. The reason is that a higher presence of YWHAG increases the probability that one of its interaction partners chooses a free YWHAG instead of increasing the size of an already existing larger complex around YWHAG.

#### Changes in singleton fractions

We also examined the influence of perturbations on the singleton fraction of each protein. Figure 12 left shows the relation between the number of interactors and the singleton fraction of a protein in the constrained network. More interactors mean more interaction possibilities, which lead to more bound copies and thus fewer singleton copies of a protein. The right side of Figure 12 shows the distribution of singleton fractions for the constrained network versus the unconstrained network. Without constraints, more interactions are possible, and therefore singleton fractions are lower overall than with constraints.

**Figure 12.**
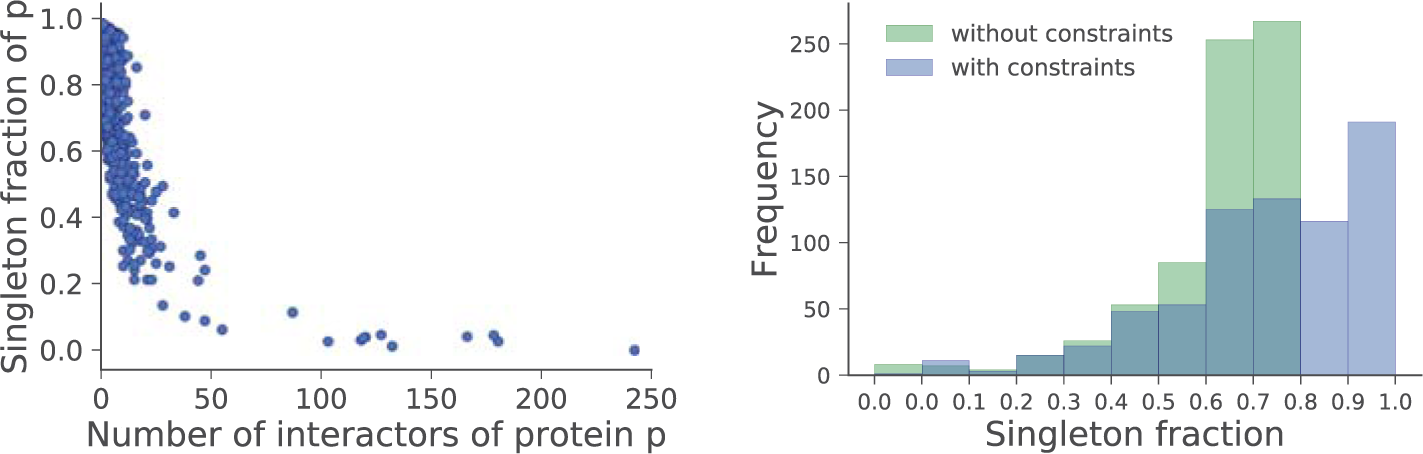
Left: Relation between number of interactors and singleton fraction for each protein. The values are averaged over 100 runs with constraints. Right: Influence of constraints on the singleton fraction distribution (averages over 100 runs with constraints and 100 runs without constraints).

We examined the average singleton fraction of each protein in unperturbed simulations as well as with overexpression and knockout of specific proteins (CRK, YWHAG, ABAT) in comparison to the unperturbed experiment. In Figure 13, the difference in singleton fraction is shown for each protein for overexpression (y-axis) and knockout (x-axis) of CRK, YWHAG and ABAT separately. We may expect that direct interactors of the perturbed protein (purple dots) are in the bottom right quadrant (more singletons after knockout and fewer singletons after overexpression of direct interactor) and that most of the other dots appear near the center with only small changes.

**Figure 13.**
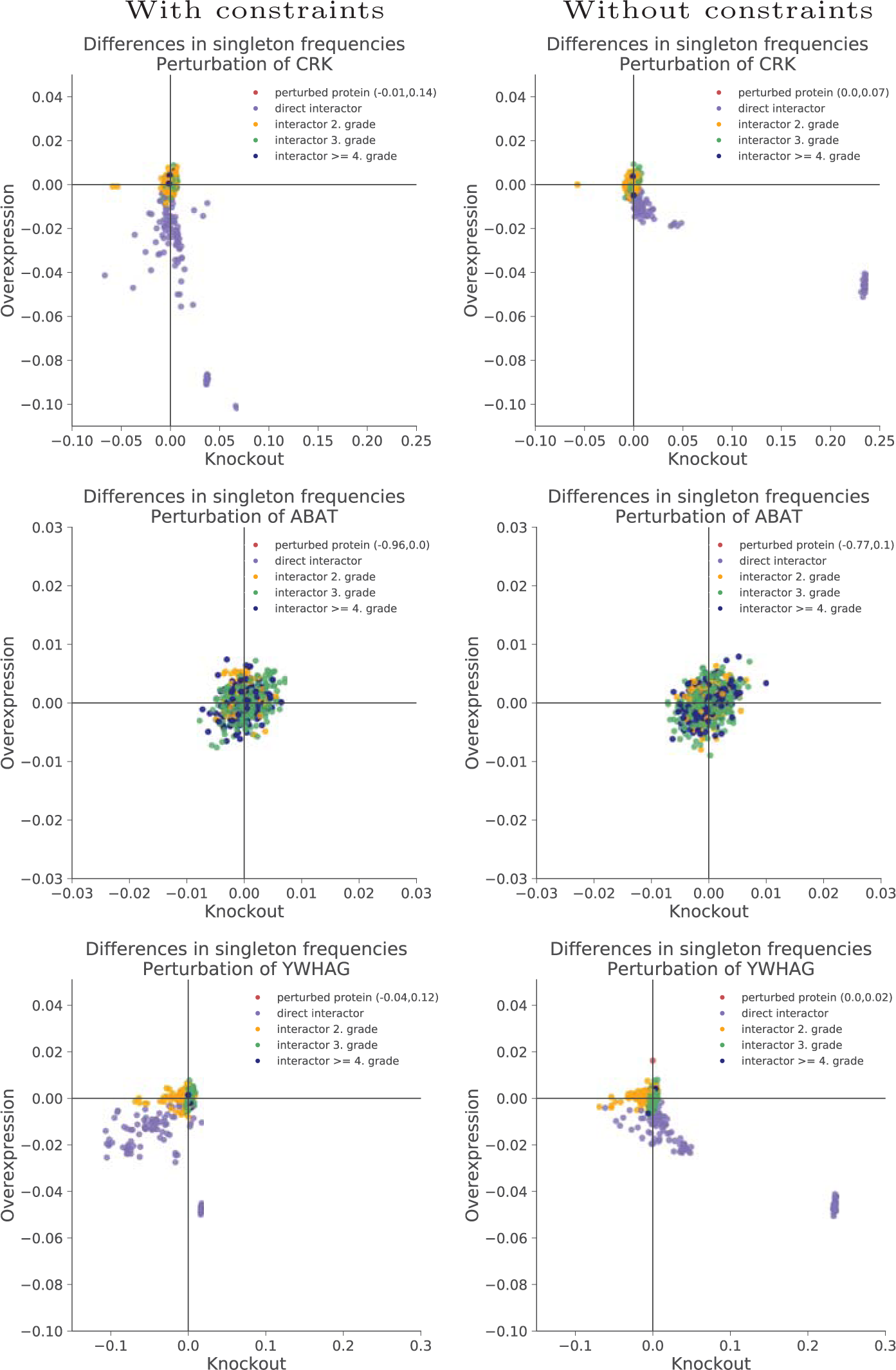
Differences in singleton fractions for perturbations (x-axis: knockout vs. unperturbed; y-axis: overexpression vs. unperturbed) of CRK (top row), ABAT (middle row) and YWHAG (bottom row) in simulations with constraints (left column) and without constraints (right column), averaged over 50 runs. Each dot represents one protein; colors show the distance (shortest path) to the perturbed protein in the interaction network. Since the perturbed protein (red dot) is not always visible, its values are given in the legend. Direct interactors (purple) are mainly expected in the lower right quadrant (more singletons after knockout, fewer singletons after overexpression). Note the different scales for the different proteins.

As expected, perturbations of ABAT have no strong effect for all proteins regardless of the distance to ABAT.

For perturbations of CRK, we observe the expected result in the network without constraints. However, in the constrained network, several direct interactors can be seen in the bottom left quadrant, meaning that those proteins have *fewer* singletons after knockout of CRK. A possible explanation is that some direct interactors of CRK may also interact with each other. In the presence of CRK, its direct interactors compete with each other. Once CRK is gone, they are free to form complexes with other proteins, especially other direct interactors of CRK. Importantly, this effect is not seen without considering interaction dependencies: Without constraints, purple dots are exclusively observed in the lower right quadrant and near the center.

Perturbations of YWHAG have a similar, but overall stronger effect than perturbations of CRK. In the unconstrained network, the majority of direct interactors can be found in the bottom right quadrant as expected, but in the constrained network, the majority shifts to the bottom left quadrant.

In summary, as CRK and YWHAG illustrate, consideration of constraints yields qualitatively and quantitatively different effects than considering the plain interaction network. Additionally, perturbations of numerically comparable proteins may lead to different results under constraints because of the local network topology: Considering interaction dependencies only of the perturbed protein and its immediate interactors may still be insufficient for foreseeing the outcome of the perturbation. Therefore, a simulation of the complete system, as our approach performs, is essential to ensure all the interactions, interaction dependencies and topological features are taken into account.

## Discussion

We have proposed a simple but powerful framework based on propositional logic for formalizing dependencies between protein interactions. As far as we are aware, this is the first such framework able to incorporate complex higher-order dependencies beyond direct competitions together with multiple copies per protein. We have shown that interaction dependencies or constraints have a direct effect on complex sizes, and additionally that they interact with local network topology. In fact, our simulations suggest that perturbations may have complex and hard-to-predict effects when taking constraints into account. The simulations are efficient in the sense that networks with a total of over a million protein copies can be simulated within under ten minutes to steady state. Compared to straightforward simulations, we achieved high speed-ups by using the bit vector techniques described in “An efficient algorithm for checking constraints” that transformed the running time from hours to minutes. The size of the network that can be simulated is thus primarily limited by the available random access memory (see table 2).

In its current form, our model makes several simplifying assumptions. For example, we check constraints only during the association phase but not during the dissociation phase. This decision is never a problem with competitions (most of the constraints in the model), and as we argue in “Simulation of protein complex formation” using the Vinculin/Talin example, we think that it captures important real effects. However, for other examples, reality might be different. By having two sets of constraints, one that has to be maintained during dissociation and one that does not, the model can be easily adapted.

Currently, our model ignores the spatial distribution of the proteins, and we have worked with uniform protein copy numbers (or concentrations) as well as uniform association and dissociation probabilities (corresponding to kinetic coefficients). Clearly, this is not realistic if the goal is to completely simulate the real biological processes happening within a cell. However, such a simulation would require much more knowledge about localization, concentrations and kinetics than is available today (late 2017). When this information becomes available, it is straightforward to scale our model accordingly. For example, association and dissociation probabilities can be chosen per interaction without causing a performance penalty and protein concentrations can already be arbitrarily parameterized in the current implementation. The spatial location of each protein copy can be considered by adding diffusion and movement rules.

Since our knowledge of constraints is currently incomplete, the biological relevance of the simulated complexes is limited. However, note that this is a problem with the available data, not with the model or simulation framework itself. Only few databases so far systematically include minable constraints, and most of them are competitions based on overlapping binding domains, as annotated in the DOMINO database [43], or data records from the IntAct database [46], which one may query with the search term `pbiorole:competitor` to obtain information on interactions where one interaction partner is a competitor. In the coming years, emerging technologies however suggest a rapid increase in the availability of the needed information, e.g., via the large scale generation of libraries of cell lines having two or more endogenously tagged fluorescent proteins [47], and recent high-throughput and multiplexed implementations of fluorescence correlation spectroscopy which allow us to systematically measure endogenous concentrations, binding constants and high-order complexes in such libraries of cell lines [48–53].

For our examples, we chose a network with a high density of known constraints. Unfortunately, the set of proteins in our network only has a small overlap with protein sets in databases of known complexes, such as CORUM [45], so we cannot directly compare predicted and real complexes. We would currently expect a number of false positive predictions, but we may also expect that the biological relevance of the simulation results will increase jointly with more complete knowledge of constraints.

We believe that our results offer important insights already today, as we demonstrated by the difference in shift of singleton fractions of direct and indirect interactors after perturbation, when comparing simulations with constraints and without constraints, but also when comparing perturbations of proteins with different network roles (with constraints); cf. Figure 13.

In the future, we will consider more realistic concentrations (or proxies for more realistic concentrations, such as setting the simulated protein copy number proportional to its number of interactors), more complete dependency data, spatial resolution, and more detailed kinetics. Moreover, we plan to extend our model to incorporate post-translational modifications such as phosphorylation, since these can also play a role in interaction dependencies. When modeling these as interactions with a special type of node, they can likewise be used within constraints.

Overall, we believe that constrained networks are a useful and versatile tool for interactomics studies that will improve and scale with increasing knowledge and data about real interaction dependencies.

## Acknowledgments

S.R. acknowledges funding from the Mercator Research Center Ruhr (MERCUR), project Pe-2013-0012 (UA Ruhr professorship) and from the German Research Foundation (DFG), Collaborative Research Center SFB 876, project C1. J.K. was supported by the Dutch NWO Veni grant 016.Veni.173.076.

